# Integrating melt-electrowriting and inkjet bioprinting for engineering structurally organized articular cartilage

**DOI:** 10.1101/2021.10.24.465623

**Authors:** A. Dufour, X. Barceló Gallostra, C. O’Keeffe, K. Eichholz, S. Von Euw, O. Garcia, D. J. Kelly

**Affiliations:** Trinity Centre for Biomedical Engineering, Trinity Biomedical Sciences Institute, Trinity College Dublin, Dublin, Ireland; Department of Mechanical and Manufacturing Engineering, School of Engineering, Trinity College Dublin, Dublin, Ireland; Johnson & Johnson 3D Printing Innovation & Customer Solutions, Johnson & Johnson Services, Inc., Irvine, CA, USA; Department of Anatomy and Regenerative Medicine, Royal College of Surgeons in Ireland, Dublin, Ireland; Advanced Materials and Bioengineering Research Centre (AMBER), Royal College of Surgeons in Ireland and Trinity College Dublin, Dublin, Ireland

**Keywords:** 3D bioprinting, MEW, Inkjet printing, Self-assembly, Spheroid, Stratified cartilage

## Abstract

Successful cartilage engineering requires the generation of biological grafts mimicking the structure, composition and mechanical behaviour of the native tissue. Here melt-electrowriting (MEW) was used to produce arrays of polymeric structures whose function was to orient the growth of cellular aggregates spontaneously generated within these structures, and to provide tensile reinforcement to the resulting tissues. Inkjeting was used to deposit defined numbers of cells into MEW structures, which self-assembled into an organized array of spheroids within hours, ultimately generating a hybrid tissue that was hyaline-like in composition. Structurally, the engineered cartilage mimicked the histotypical organization observed in skeletally immature synovial joints. This biofabrication framework was then used to generate scaled-up (50mm × 50mm) cartilage implants containing over 3,500 cellular aggregates in under 15 minutes. After 8 weeks in culture, a 50-fold increase in the compressive properties of these MEW reinforced tissues were observed, while the tensile properties were still dominated by the polymer network, resulting in a composite construct demonstrating tension-compression nonlinearity mimetic of the native tissue. Helium ion microscopy further demonstrated the development of an arcading collagen network within the engineered tissue. This hybrid bioprinting strategy provides a versatile and scalable approach to engineer cartilage biomimetic grafts for biological joint resurfacing.

## 1. INTRODUCTION

Under normal physiological conditions, articular cartilage is capable of transmitting loads of several times body weight through synovial joints for decades [1]. However cartilage injuries, if left untreated, can progress to more serious degenerative articular conditions due to the avascular nature of the tissue and its relatively limited regenerative capacity [2]. Osteoarthritis (OA) is characterized by progressive loss of articular cartilage tissue and function and is a debilitating disease affecting millions of people worldwide [3,4]. For patients suffering from end-stage OA, total joint replacement is the standard surgical treatment to restore mobility. While this procedure is well established, it does not provide a long-term solution because of the limited lifespan of the synthetic prostheses (∼ 25 years [5]), and revision surgeries are often required for a variety of reasons such as wear, loosening, or instability [6]. Hybrid synthetic-biological implants recapitulating the core structure and function of cartilage could potentially be used to treat damaged synovial joints and delay or potentially prevent the development of OA, but the engineering of such grafts remains a significant challenge.

Scaffold-free tissue engineering strategies represent a particularly promising route to the engineering of functional articular cartilage grafts [7]. Such approaches rely on a non-adherent substrate to force a high-density cell population to aggregate, with abundant cell-cell interactions mediated by N-cadherin [8]. This mimics the process of mesenchymal condensation and cartilage tissues form in a manner reminiscent of cartilage morphogenesis [8,9]. At equivalent cell-seeding densities, tissues engineered *via* such self-assembly processes show more robust and faster hyaline-like matrix accumulation over standard hydrogel encapsulation [10,11], and can generate cartilage tissues with biochemical and biomechanical values within the range of native articular cartilage [12–14]. It has also been shown that radial confinement increases collagen organization within self-assembled cartilage [15]. This is of particular interest since recapitulating the complex zonal organization of articular cartilage remains an important challenge [16], as this structure is integral to the ability of the tissue to withstand the challenging mechanical loading of synovial joints [17–19]. To further improve the structural organization of engineered cartilage tissues, hybrid approaches that combine the benefits of cellular self-assembly or self-organization with biofabrication techniques such as 3D bioprinting have recently been developed. Printed polymeric structures can be used to guide the deposition of a cartilage biomimetic collagen network within engineered tissue [20]. To date, polymeric structures with relatively thick fibers generated by fused deposition modeling (FDM) have been used to trigger cellular self-assembling and to direct subsequent tissue organization [20,21]. These large polymeric fibers lead to the development of overly stiff and non-compliant constructs [22] that are not mimetic of the native tissue and could potentially damage opposing joint surfaces if implanted *in vivo*. Ideally, polymeric reinforcement within such hybrid engineered tissues would mimic the functionality provided by the collagen network in articular cartilage, specifically a fibrillar matrix that primarily sustains tensile loads [19], but in isolation contributes little to the compressive properties of the tissue [23].

Melt-electrowriting (MEW) uses voltage-stabilized jets to accurately place low-micrometer-scale fibers in pre-defined locations in 3D space [24]. The fiber diameter ranges from 820 nm [25] to 140 μm [26]; whereas it is usually over 200 μm with FDM, limiting the capacity of this addidative manufacturing technology to produce truly biomimetic implants [27,28]. In contrast, MEW enables the development of highly porous (80-98 vol% pore volume), sophisticated and biomimetic scaffolds [29,30]. For example, MEW has been used to mimic the anisotropy of the collagen network in cartilage [31,32], mechanically reinforcing hydrogels in a way that recapitulates the behaviour of collagens in cartilage [31]. MEW scaffolds have also been used as a substrate for the assembly of pre-formed multicellular spheroids [33,34]. Here, we hypothesized that a MEW network could guide cartilage-specific tissue organization during the growth of self-assembled cell aggregates, while simultaneously reinforcing the resulting hybrid construct in a manner analogous to that of the collagen network in articular cartilage, specifically providing tensile strength and stiffness but contributing little to the compressive properties of the tissue in the absence of proteoclycans. To this end, we combined MEW and inkjet bioprinting into a sequential biofabrication framework where a defined number of mesenchymal stem cells (MSCs) were ink-jetted into box-like MEW scaffolds, which supported spontaneous cellular aggregation within each microwell (**Figure 1**). We demonstrate how the association of such additive manufacturing technologies can be used to produce sheet-like tissue constructs composed of multicellular spheroids maturing into stratified cartilage tissue, within only a very limited fraction of synthetic polymer (< 2%). By expanding the capabilities of this novel multiple-tool biofabrication strategy, we also demonstrate that it can be used to engineer large and functional cartilage grafts with potential applications in biological joint resurfacing.

**FIGURE 1:**
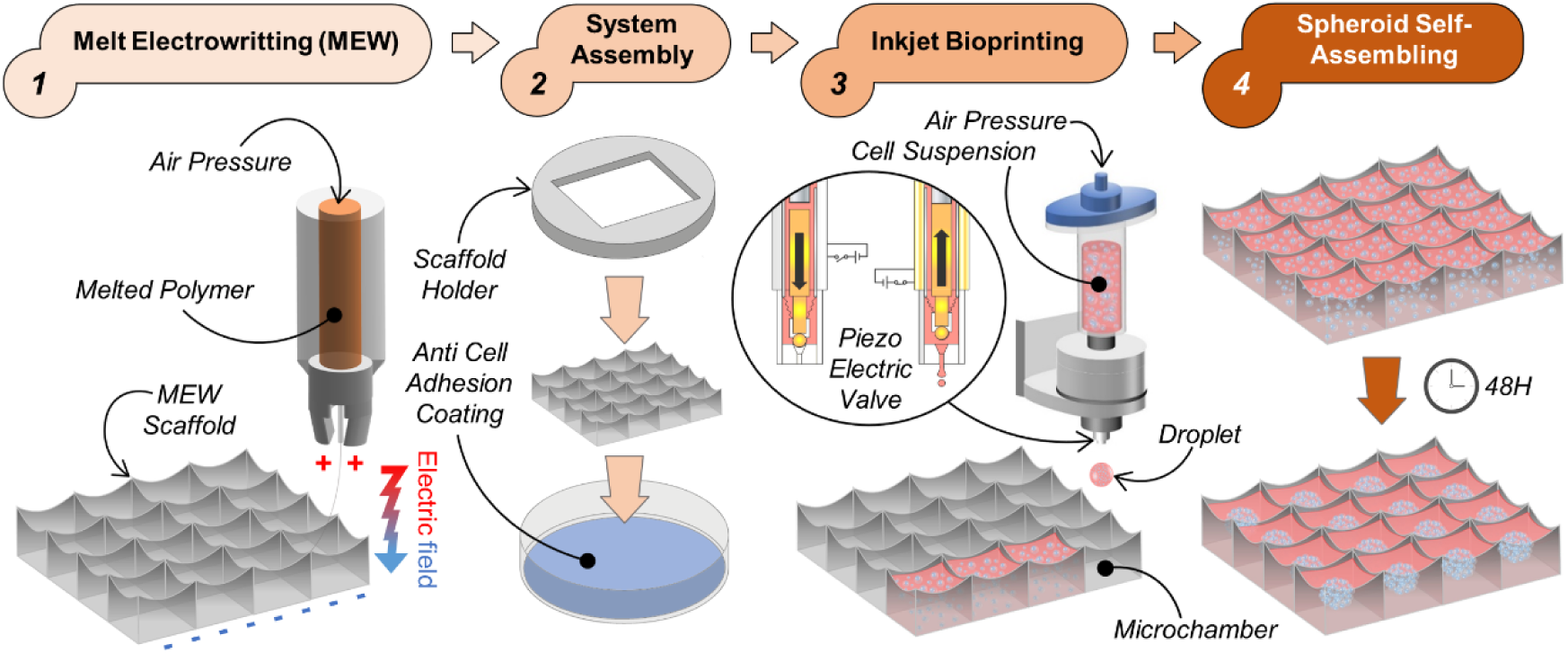
Biofabrication process. A box-like structure made of fibers in the micron-size range is produced by extruding poly(ε-caprolactone) (PCL) across an electric field (*Melt-Electrowriting* or MEW). The MEW scaffold is centered in a plastic dish coated with a solution preventing cell adhesion to the dish (*System Assembly*). A droplet containing a defined number of cells is then printed in every single chamber of the scaffold through a piezoelectric valve (*Inkjet Bioprinting*). As cell adhesion is limited at the bottom of the assembly by the hydrophobic coating, and on the sides by the hydrophobic polymer, the cells aggregate and self-assemble into spheroids within the MEW scaffolds within 48H (*Spheroid Self-Assembling*).

## 2. RESULTS

### 2.1. Integrating melt-electrowriting and inkjet printing to generate arrays of cellular spheroids

The first step of the present work was to elaborate a strategy to trigger the self-assembly of cellular aggregates from cells ink-jetted into micron-sized polymeric microchamber systems (**Figure 1**). To this end, melted polycaprolactone (PCL) was extruded across an electric field to produce an orthogonal array of microfibers (≈ 7 μm diameter) (**Supplemental Table 1**) (**Supplemental Figure 1.A-D**). The microchamber height (≈ 0.75 mm) and spacing (≈ 0.8 mm) were kept constant throughout, resulting in a microchamber volume of 0.48 mm^3^. Subsequently, the MEW scaffold was placed onto a non-cell adhesive dish coated with poly(2-hydroxyethyl methacrylate) (poly-HEMA) that defined the temporary bottom boundaries and supported cell-aggregation post inkjetting. Lastly, the printed microchambers were loaded with cells by inkjet printing a cell suspension into each microchamber. The valve opening time, which defines the volume of one drop printed through a single valve opening, was kept constant and reproducible volumes were printed throughout the experiments (**Figure 2.A**). After identifying the volume corresponding to a single drop/valve opening (1.16 ± 0.06 μl), the concentration of the cell suspension was defined (30 million cells/ml) and inkjet printing was used to seed a defined number of cells suspended in cell culture medium into the confining hydrophobic polymeric microchambers (34,845 ± 1,744 cells per microwell) (**Figure 2.B**).

**FIGURE 2:**
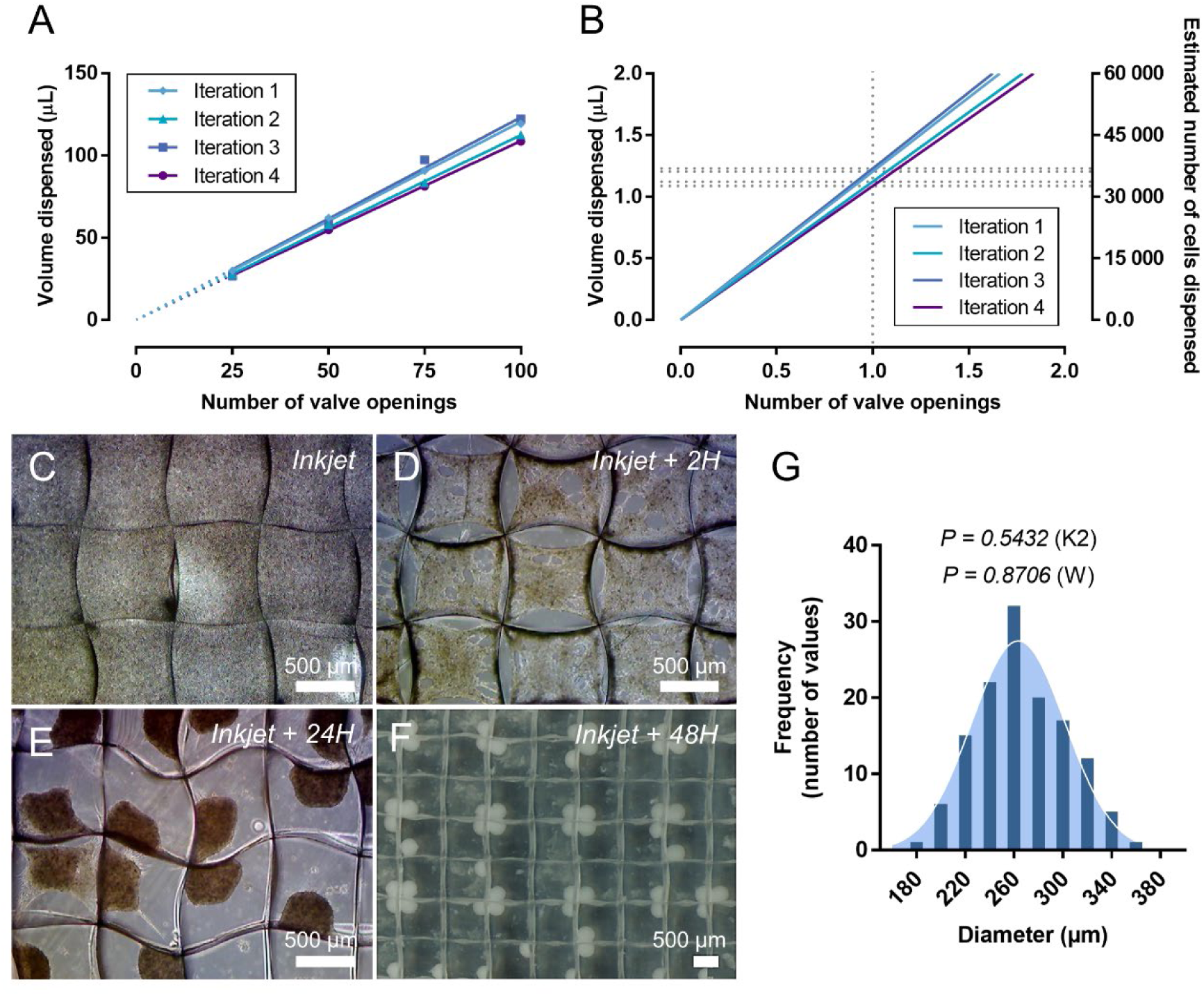
Characterisation of inkjet bioprinting and self-assembly processes. To characterize the inkjet process, (A) a curve displaying the volume dispensed against the number of valve openings is generated for constant valve opening time and pressure settings and extrapolated to zero (dotted lines). (B) The number of cells dispensed during a single valve opening is estimated from this extrapolation and for a defined cell concentration. (C-D) Cells in box-structured MEW scaffolds (C) immediately, (D) 2H, and (E) 24H after inkjet. (F) Macroscopic picture of the self-assembled spheroids 48H after inkjet bioprinting. (G) Frequency distribution analysis of the spheroid diameter (dark blue bars) fitted with a Gaussian curve (light blue bell-like motif). P-values of d’Agostino-Pearson (K2) and Shapiro-Wilk (W) normality tests are indicated for α = 0.05.

Microscopic observations made immediately after printing showed a homogeneous cell suspension segmented by protruding MEW fibers, indicating individual filling of the microchambers with cells (**Figure 2.C**). A contracting cell layer was observed in each microchambers a few hours later (**Figure 2.D**), further condensing with time (**Figure 2.E**), and resulting in a structured array of cellular spheroids within the MEW template 48 hours later (**Figure 2.F**). Interestingly, spheroids self-assembled predominantly in the corner of the microchambers and against each other, exhibiting a regular pattern. Temporal monitoring of the self-assembling process showed that a critical point is reached during the contraction phase where the spheroid detaches almost totally from the scaffold, keeping just one or two points of attachment that hauled the spheroid to a specific corner (**Supplemental Video 1**). Closer examination confirmed spheroids nesting at the bottom of the polymeric chambers (**Figure 3.A, B**), with signs of attachment to the microfibers (**Figure 3.B-D**) and cellular extensions protruding through the fiber walls (**Figure 3.D**). This last observation suggests that the spheroids could be able to communicate physically through the fiber wall, facilitating their self-assembling into a regular pattern and eventual fusion. Finally, the spheroid diameter (266 ± 37 μm) was shown to be normally distributed (**Figure 2.G**), indicating that spheroids of reproducible size can be generated in every single microchamber of the MEW scaffold. Taken together, these results demonstrate that cellular condensation occurred following deposition of the cell suspension into microwells and that structured arrays of cellular spheroids can be engineered by integrating inkjet printing and MEW.

**FIGURE 3:**
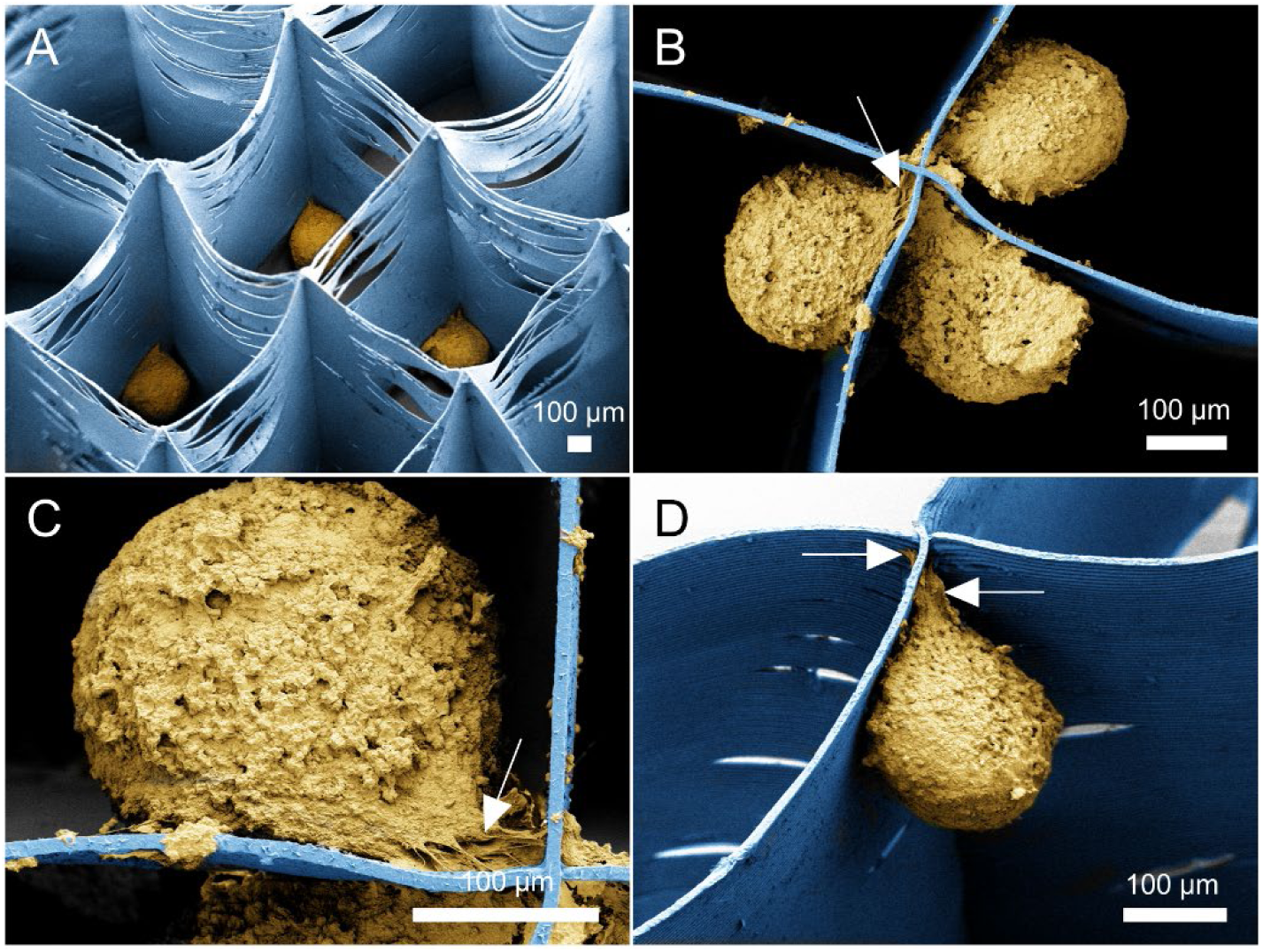
Scanning electron microscopy (SEM) images of self-assembled spheroids in MEW scaffolds 48 hours after inkjet printing. The scaffolds were imaged from (A) the top or (B-D) the bottom. (A) The top view image shows spheroids sitting in the corners of MEW microchambers. Bottom views show (B) spheroids that associate against the wall of fibers and (C, D) attachment to the fibers (indicated by white arrows). The MEW scaffold is colored in blue and cell spheroids in yellow. Please note: spheroid shrinkage occurred in preparation for SEM.

### 2.2. Self-organization of hyaline-like cartilage in MEW scaffolds following inkjet bioprinting

The structured array of cellular spheroids formed within the PCL template grew over time to fill the microchambers, with fusion between adjacent spheroids evident after 21 days macroscopically and in live/dead imaging (**Figure 4**). The isolated spheroids observed at day 0 grew out of their chambers to fuse with their neighbours (**Figure 4.C**) and covered the surface with viable cells (**Figure 4.E, F**). Furthermore, microfiber walls clearly visible at day 0 and in empty scaffolds (**Figure 4.B**) could not be distinguished on hybrid tissue cross-sections that show a continuous tissue with a glossy appearance similar to native cartilage. Histological staining for sGAG deposition confirmed robust cartilage development and the formation of a highly connected material (**Figure 5.A**), with the sGAG content of the engineered tissues (2.5 ± 1.2 ww%) approaching that of native articular cartilage (∼ 3-10 ww% [18,35]) (**Figure 5.B**). Although total collagen content of engineered tissues (0.9 ± 0.2 ww%) was an order of magnitude below that of articular cartilage (∼ 5-30 ww% [18,35]), hybrid tissues were hyaline-like in composition as evidenced by strong positive staining for type II collagen while type I collagen was barely detected (**Figure 5.C**).

**FIGURE 4:**
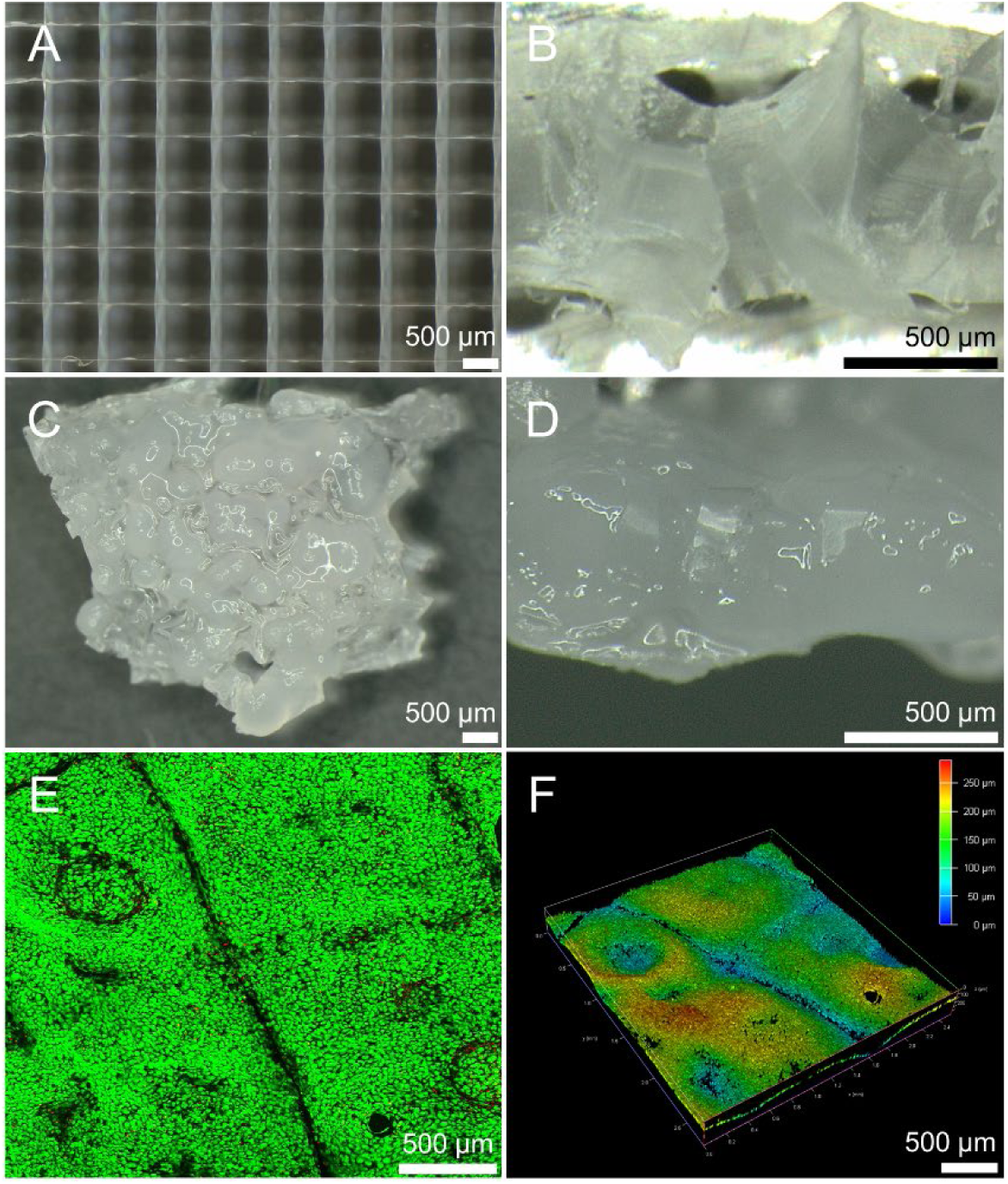
Tissue formation following 21 days of chondrogenic induction. Macroscopic images of (A, B) empty and (C, D) cell-printed scaffolds. Right pictures (B, D) are cross-section views. (E-F) Live dead staining of cells in the MEW scaffolds (showing live cells in green and dead cells in red) with (F) depth reconstructions of the green (live) channel.

**FIGURE 5:**
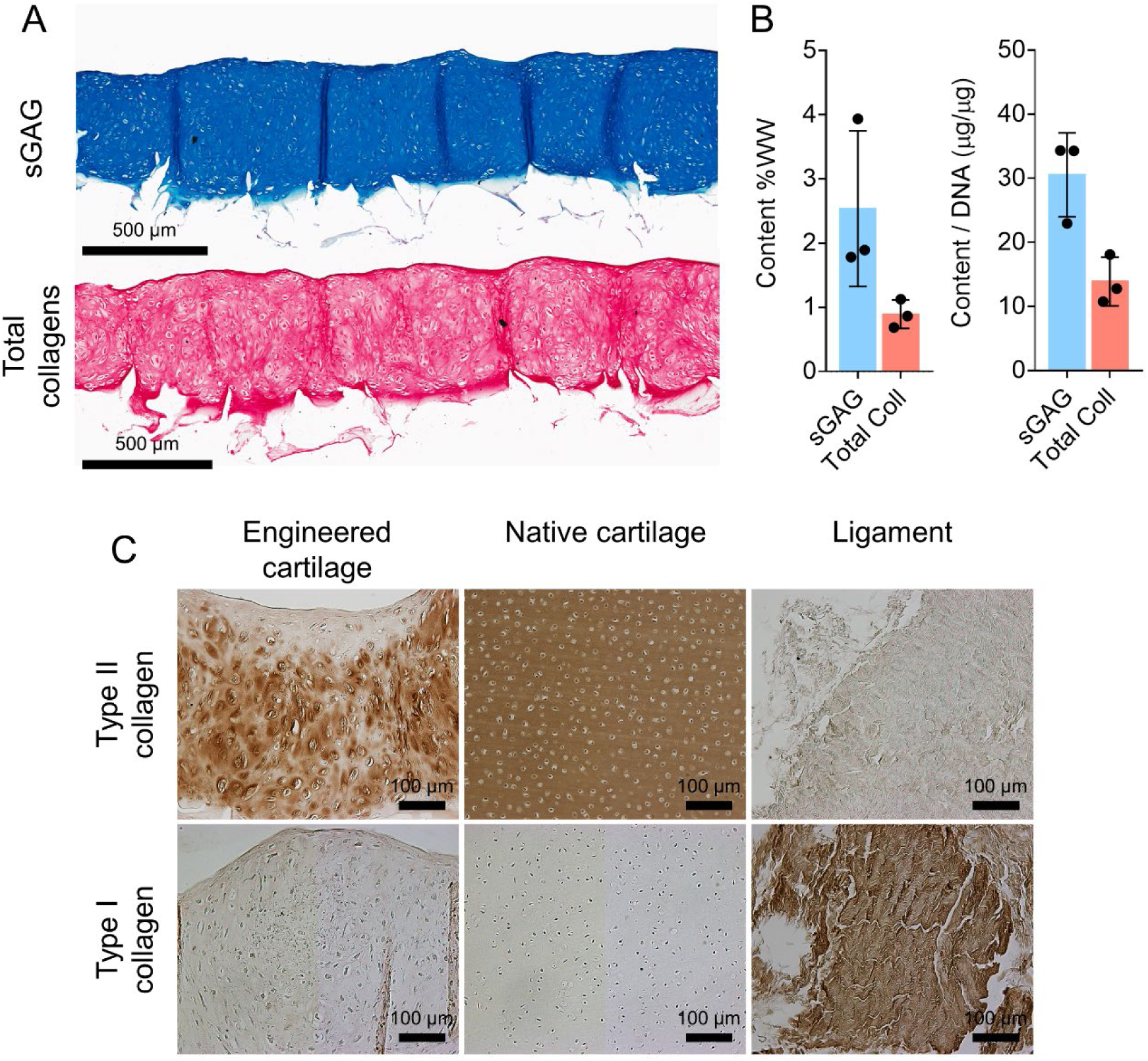
Histological analysis and biochemical content of the engineerd tissue after 21 days of culture. (A) Parallel tissue sections stained with alcian blue (sGAG) and picrosirius red (total collagen). (B) Quantitative evaluation of the sGAG and total collagen contents in the biofabricated tissues normalized to wet weight (WW) or DNA content. The values given in the dot plots represent scaffolds as data points, mean as a bar, and standard deviation as error bars (n = 3). (C) Immunohistochemistry for type I and type II collagen with native cartilage and ligament as positive or negative controls.

### 2.3. Engineered cartilage within MEW scaffolds mimics the spatial organization of articular cartilage from skeletally maturing joints

Microscopic observations of hybrid tissue sections stained for hematoxylin-eosin revealed cells with round morphology typical of chondrocytes which were randomly distributed within the tissue but flattening parallel to the surface in the superficial layers (**Figure 6.A**). A similar cell organization is found in developing cartilage and is known to be associated with an organized collagen network. Therefore we next used polarized light microscopy (PLM) to determine the degree of organization of collagen fibrils in the engineered tissues, and compared them to articular cartilage from skeletally immature joints (**Figure 6.B-E**). Tissue sections from native cartilage displayed multi-zonal features typical of maturing cartilage [18,36,37] (**Figure 6.B**). A greenish mildly birefringent radial zone with collagen fibrils oriented perpendicular to the surface is sandwiched between a tangential zone at the surface and an isotropic zone underneath (where collagen fibre reorganization occurs), both showing a strong yellow birefringence with collagen fibre pattern-oriented parallel to the surface. A very similar collagen architecture was found in the engineered tissues, with a 3 layers organization matching that found in maturing tissue. Remarkably, the thickness of the radial zone between tissues is similar, as was the architeture of the collagen fibrils under polarized light (oriented at ∼ 80-70°). The average fibril orientation and dispersion were then assessed in the tangential, radial, and isotropic zones of the tissues. Average fiber orientation was 4.55 ± 4.61°, 80.97 ± 7.36° and 14.49 ± 13.86° in the engineered tissue and 1.5 ± 1.2°, 85.89 ± 3.35° and 5.17 ± 4.96° in native maturing cartilage (**Figure 6.C**). Similarly, fiber dispersion was 18.71 ± 6.4°, 16.82 ± 6.47° and 19.31 ± 10.12° in engineered tissue and 8.49 ± 2.76°, 16.84 ± 2.42° and 16.01 ± 1.93° in native cartilage (**Figure 6.D**). Statistical analyses revealed little differences between the engineered and native tissues, highlighting the similarity in their collagen organizations. Coherency was also used as a measure of the anisotropy, tending to 1 if there is a dominant direction in the average region. Coherency differed only between radial zones, where a higher degree of organization was found in the engineered tissue (0.5 ± 0.09) compared to juvenile cartilage (0.24 ± 0.07) (**Figure 6.E**). Taken together, this analyses demonstrate spatial changes in cell morphology and the organization of collagen fibrils through the depth of the engineered tissue that mimicked that seen in the articular cartilage of skeletally immature joints.

**FIGURE 6:**
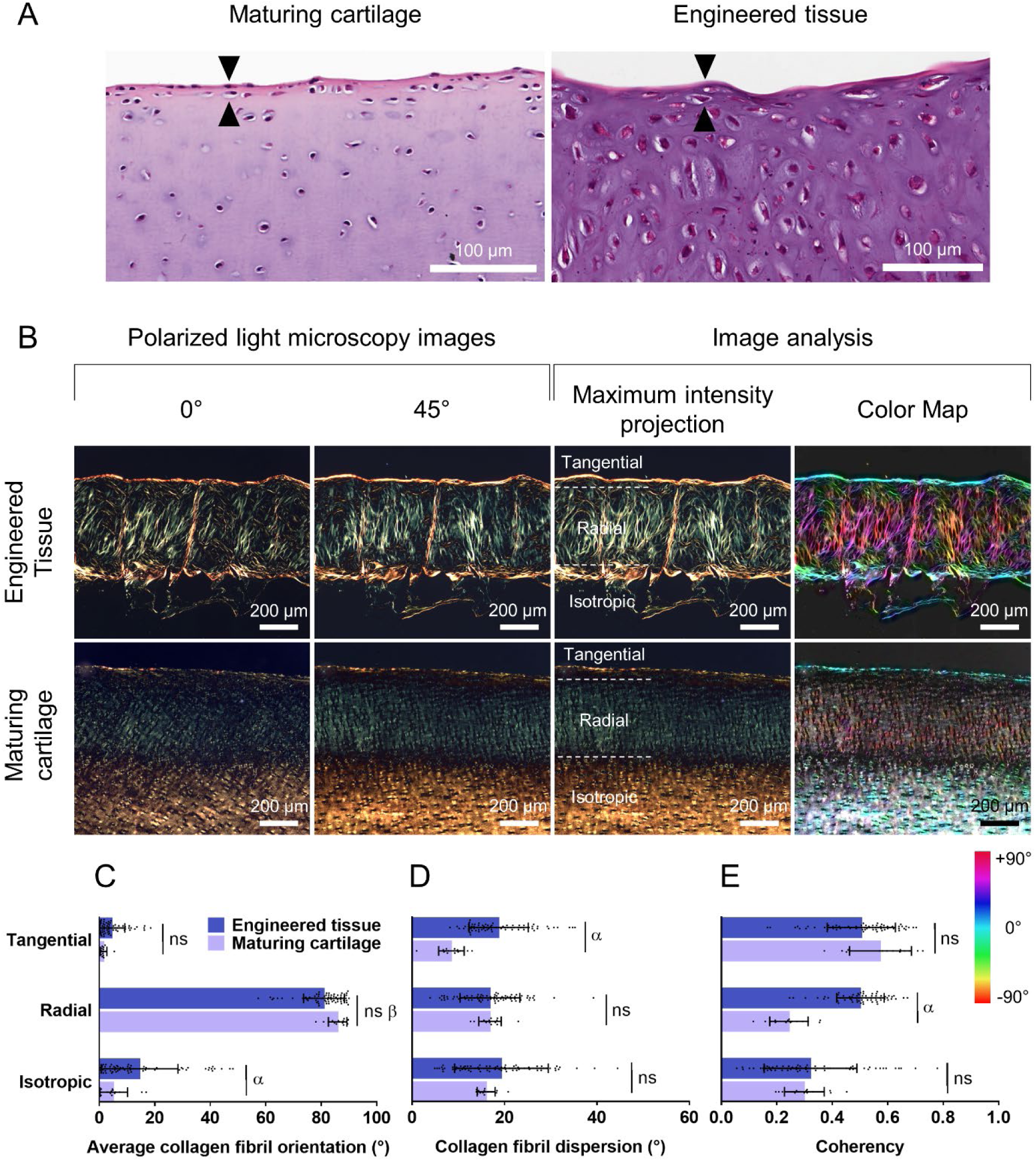
Engineering of structurally organized articular cartilage. (A) High magnification pictures of native (maturing) and engineered tissue sections stained for hematoxylin and eosin. The black arrows highlight cells flattening in the superficial layer. The middle panel (B) shows representative polarized light microscopy (PLM) images (x10 magnification) for two different angles (0 and 45 degrees) that were merged (maximum intensity projection) and analyzed with ImageJ plugin OrientationJ (color map). (C, D, E) Quantification of collagen fibril (C) orientation, (D) dispersion, and (E) coherency in the different zones observed in merged PLM images. Coherency is a measure of local orientation and isotropic properties, ranging from 0 if the image is isotropic in the analyzed region of interest to 1 when the local structure has one dominant orientation. The values given in the dot plots represent x10 histological regions as data points, mean as bar, and standard deviation as error bars. Multiple regions analyzed per sample (n = 3). ns indicates no statistically significant difference between maturing and engineered tissues for the indicated zone (P > 0.05). α indicates a statistically significant difference between maturing and engineered tissues for the corresponding region (P < 0.01). β denotes significance compared to tangential or isotropic zone, regardless of the nature of the tissue (native or engineered) (P < 0.001) (one-way ANOVA).

### 2.4. Scaled-up engineered cartilage possesses mechanical properties approaching that of native articular cartilage

We next explored whether our multi-tool biofabrication strategy could be used to produce implants of a size suitable for resurfacing complex articular surfaces such as the hip joint (**Supplemental Figure 1.E-F**). To do this, 60 × 60 mm MEW scaffolds were printed and maintained in a poly-HEMA coated 60 mm Petri dish with a moulded PCL holder so that the inkjet area was a 50 × 50 mm square. Cells were inkjet printed in the 3,591 microchambers in this region in about 15 minutes without interruption. A cell spheroid was obtained in every pore where cells were ink-jetted, with microtissues developing and merging only into the ink-jetted area over 8 weeks and leaving the edge of the scaffold empty (**Figure 7.A and B**). To evaluate if the engineered hybrid tissue was functional, a combined stress-relaxation and dynamic unconfined compression protocol was used to determine the mechanical properties of the tissue. Tissue constructs stress-relaxed and strain-stiffened similarly to articular cartilage [38,39] (**Figure 7.C**). The compressive modulus of engineered tissues was 177 ± 30 KPa and 388 ± 55 KPa at 20% and 30% strain respectively (**Figure 7.D**), which represents a 50-76 fold increase compared to MEW scaffolds (4 ± 2 KPa and 5 ± 3 KPa) and approached native tissue properties (0.24 to 1.4 MPa [39–41]). Equilibrium and dynamic modulus are two other important parameters used to quantify the mechanical function of engineered cartilage tissues [42]. The equilibrium modulus is a measure of the compressive stiffness of the tissue solid matrix since it is recorded after the ramp and hold phase when fluid is no longer moving through the tissue. The equilibrium modulus of engineered composite tissues was 180 ± 13 KPa and 214 ± 24 KPa at 20% and 30% strain respectively (**Figure 7.E**), which is close to that of native tissue (0.2 to 2 MPa [18,39,43–46]) and represents a 20-27 fold increase compared to MEW scaffolds (9 ± 2 KPa and 8 ± 2 KPa). Lastly, during the dynamic phase of the test, a cyclic displacement is applied to the tissue to test its capacity to generate fluid pressurization / load support, which is related to the permeability of the solid matrix. Dynamic moduli recorded for tissue constructs were 1,4 ± 0,2 MPa at 20% strain and 2,6 ± 0,3 MPa at 30% strain (**Figure 7.F**). These values are of a similar order of magnitude as the native tissue (5 to 60 MPa [39,47– 51]) and represent a marked 26-51 fold increase compared to empty scaffolds (56 ± 18 KPa and 51 ± 23 KPa).

**FIGURE 7:**
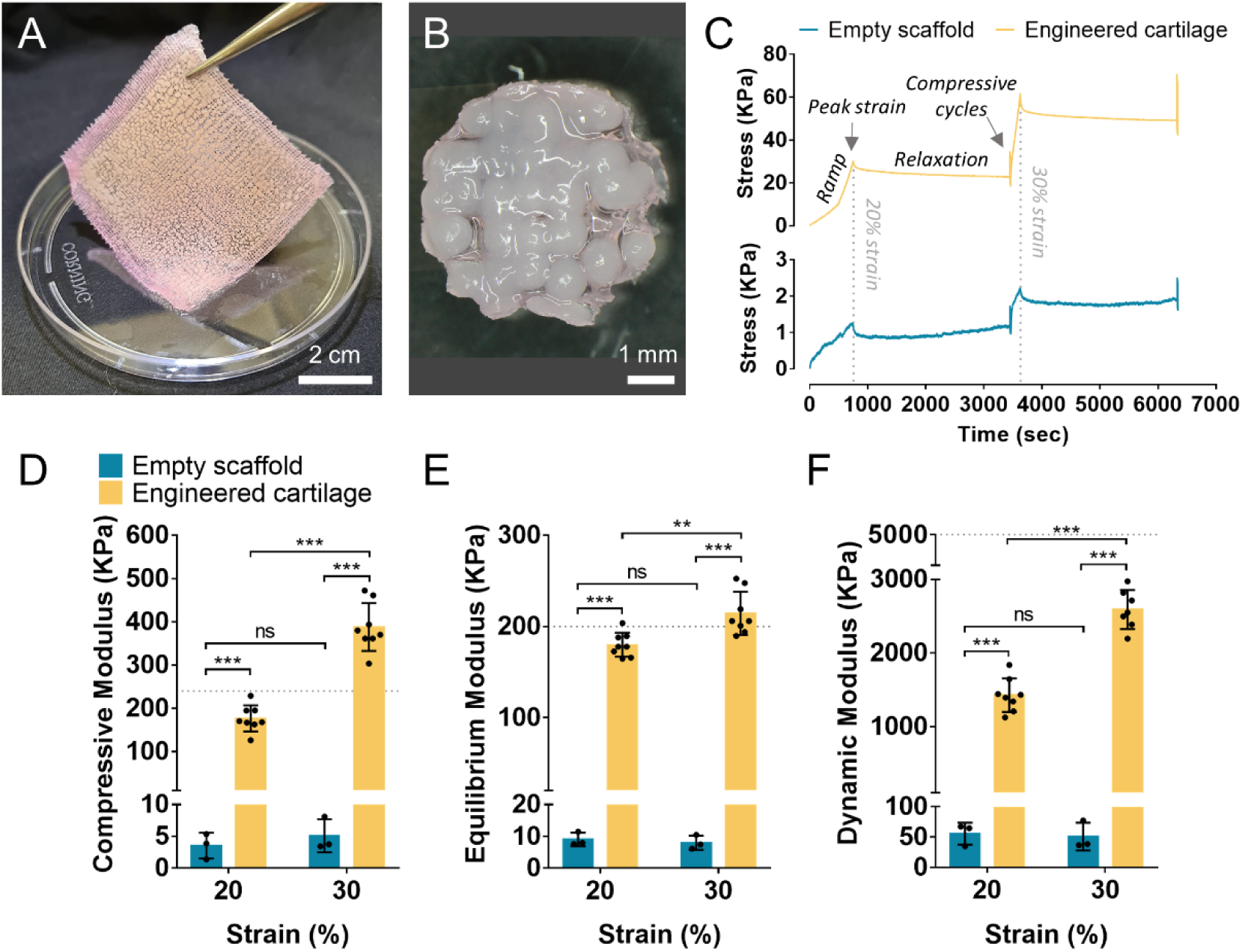
Integrating melt-electrowriting and inkjet for engineering large cartilage graft. MEW scaffolds of 60 × 60 mm were printed and bone marrow-derived MSCs were ink-jetted into all microchambers covering a square surface of 50 × 50 mm, then cultured for 8 weeks in chondrogenic media. Two scaffolds were produced. (A) Large MEW scaffold showing complete filling of the ink-jetted area with self-assembled spheroids. The scaffold is handled above the lid of a 60 mm cell culture petri dish demonstrating ease of handling. (B) Five-millimeter diameter cylindrical construct punched out of the main scaffold for unconfined compression testing. (C) Representative stress-time curve of tissue-engineered cartilage (yellow line) and empty MEW scaffold (blue line) highlighting the different steps of the unconfined compression testing procedure. (D) Compressive (or ramp) modulus, (E) equilibrium modulus, and (F) dynamic modulus in unconfined compression of engineered tissue (yellow bars) and empty scaffold (blue bars) when applying increasing levels of strain amplitude. The values given in the dot plots represent punched-out regions as data points, mean as a bar, and standard deviation as error bars. ** and *** indicate statistically significant differences (**p < 0.01, *** p < 0.001), whereas “ns” denotes no significance (one-way ANOVA). Dotted lines indicate the minimal value recorded for articular cartilage in literature.

The tensile properties of the larger, scaled-up tissue-engineered cartilage composites were also investigated (**Figure 8.C-G**). Scaffolds were successively stretched and relaxed at 3, 6, and 9 % strain, then ramped beyond the yield point (**Figure 8.D**). The overlapping of the stress-strain curves first suggested minimal differences in tensile properties between engineered cartilage and the empty MEW scaffolds. At 9% strain, tensile Young’s modulus was 1.4 ± 0.3 MPa and 0.9 ± 0.2 MPa (**Figure 8.E**), and equilibrium modulus was 1.7 ± 0.1 MPa and 1.5 ± 0.1 MPa (**Figure 8.F**) for engineered tissues and empty scaffolds respectively. These values approach that of native cartilage (5-12 MPa for tensile Young’s modulus [41] and 5-25 MPa for equilibrium modulus [43,52,53]), but no significant differences were found between composite tissues and empty scaffolds. However, only the engineered cartilage composite displayed a strain-stiffening behaviour. Although it remains significantly lower than that of native tissue (0.8-25 MPa [54]), higher yield stress was recorded for engineered tissues (132 ± 29 KPa) compared to MEW scaffolds (96 ± 14 KPa), pointing to some reinforcement in tensile properties after tissue maturation. Taken together, these results demonstrate that upscaling the hybrid biofabrication process results in functional analog tissues that could potentially be used for joint resurfacing.

**FIGURE 8:**
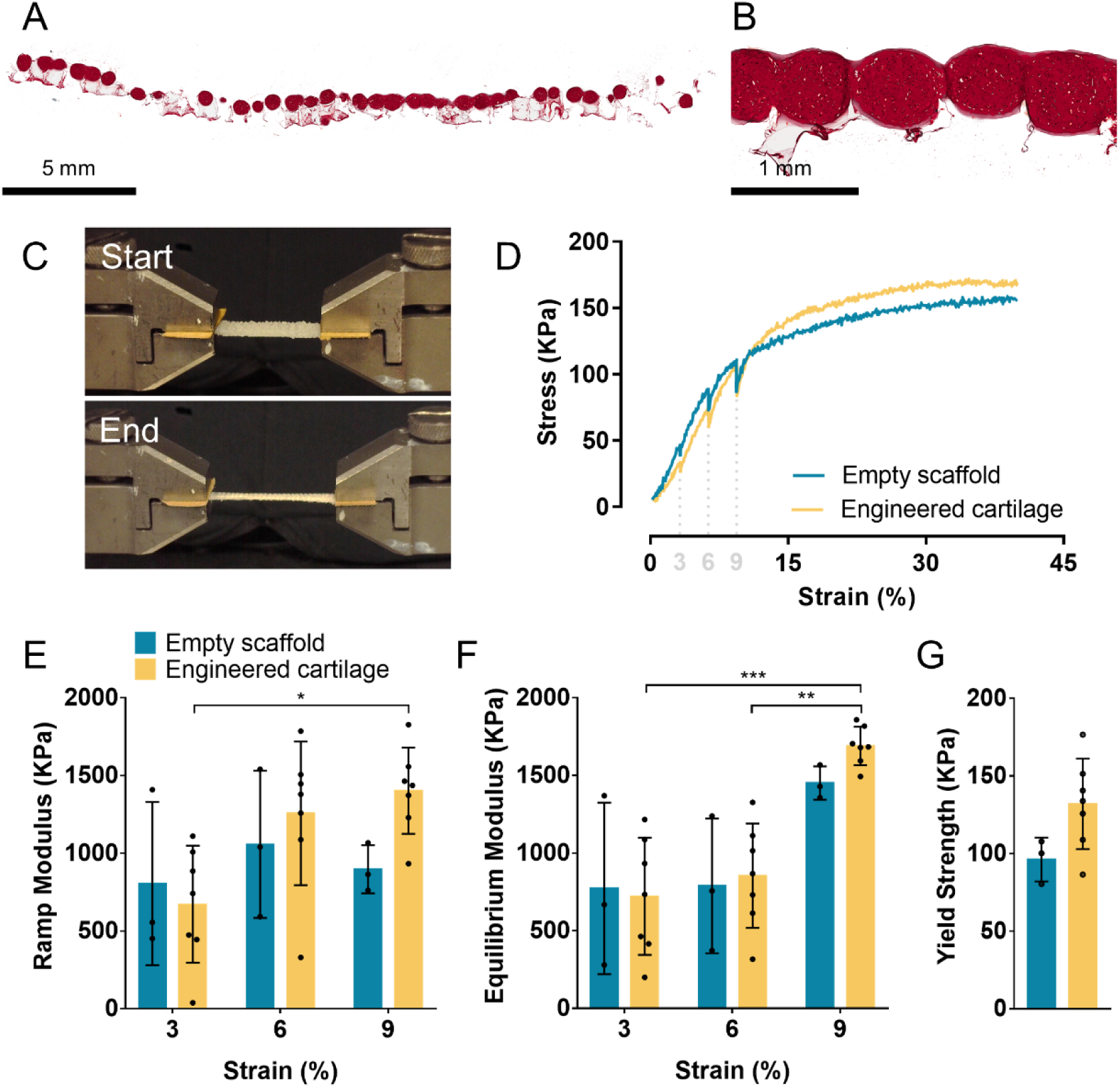
Uniaxial stress-relaxation tensile testing of large engineered tissues. (A, B) Histological cross-section of engineered tissue stained for safranin-O and observed at (A) very low and (B) low magnification. (C) Pictures of a 20 × 5 mm tissue section sampled from the engineered tissue at the start and the end of the tensile testing procedure. (D) Representative stress-strain curve of tissue-engineered cartilage (yellow line) and empty MEW scaffold (blue line). Dotted lines indicate peak strain at 3, 6, and 9% strain that were each followed by relaxation before increasing strain amplitude. (E) Ramp and (F) equilibrium tensile modulus as well as (G) yield strength in uniaxial stress-relaxation tensile testing when applying increasing levels of strain amplitude on engineered tissue (yellow bars) or empty scaffold (blue bars). The values given in the dot plots represent test sections sampled from the engineered tissues or empty scaffolds as data points, mean as a bar, and standard deviation as error bars. *,** and *** indicate statistically significant differences (*p < 0.5, **p < 0.01 and *** p < 0.001, one-way ANOVA).

Lastly, helium ion microscopy (HIM) was used to visualize in more detail the spatial organization of the collagen fibrils in the engineered tissue (**Figure 9**). Clear spatial changes in structural organization with depth were observed (**Figure 9.B1 and B2**). Collagen fibrils had a predominantly parallel orientation at the surface of the tissue (**Figure 9.C1, F1, and C2, F2**), and arcaded (**Figure 9.D1, G1, and D2, G2**) to a perpendicular orientation more deeply (**Figure 9.E1, H1, and E2, H2**), thus resulting in a Benninghoff-like architecture. Small fibrils (38 ± 12 nm in diameter) with abundant fibrillar connections were observed through the tissue, similar to maturing articular cartilage. The collagen network was denser toward the surface of the tissue with packed fibrils, a feature observed in mature tissue [18]. These findings demonstrate that the hybrid biofabrication process can generate tissue analogs with spatial changes in collagen fibril orientation mimicking that of native cartilage.

**FIGURE 9:**
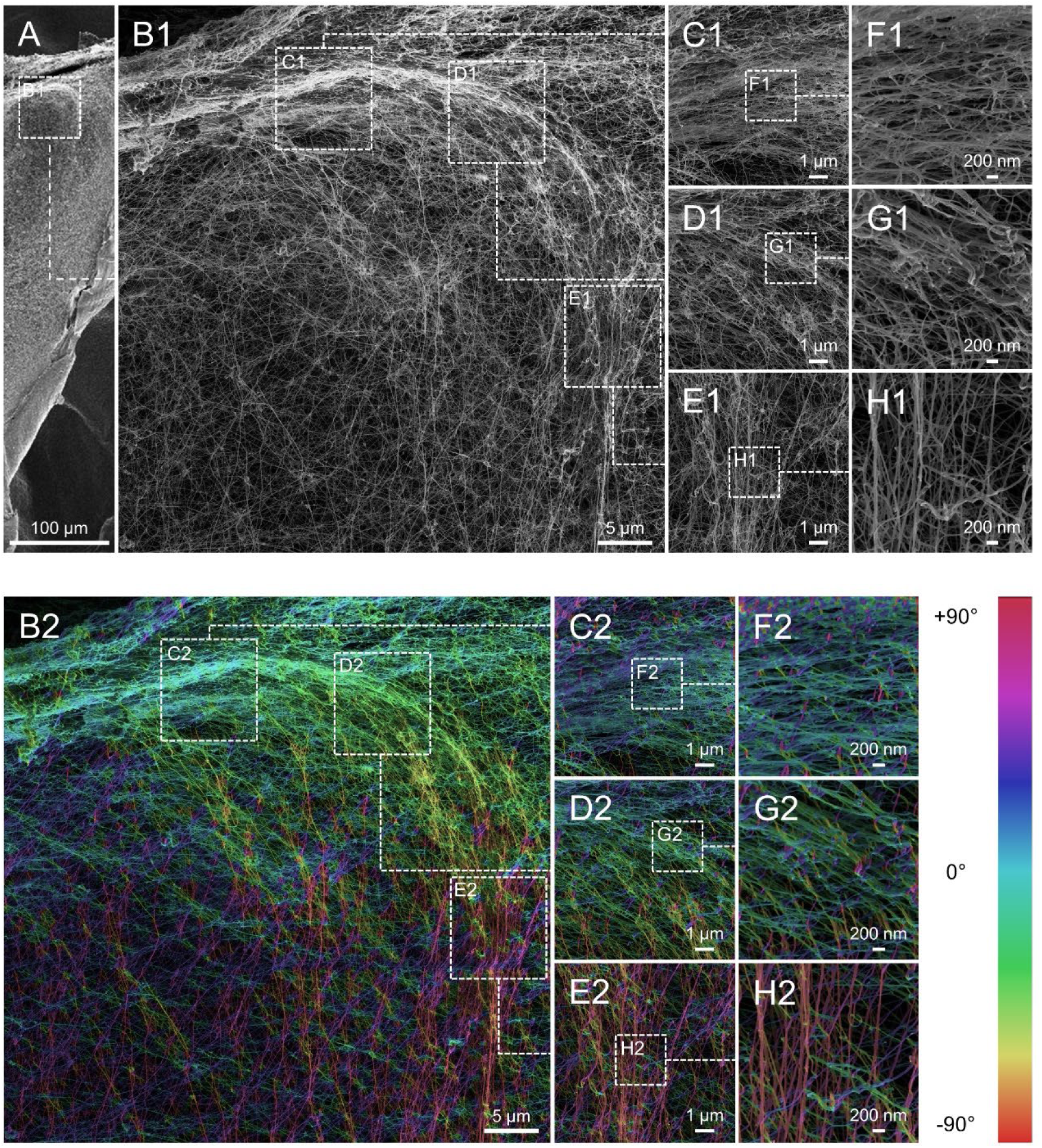
High-resolution visualization of collagen fibril orientation. The upper panel (A, B1-H1) shows helium ion microscopic images of the collagen network within engineered tissue cultured for 8 weeks in chondrogenic media. High-resolution images were acquired at the surface of the sample cross-section to observe the shift in fibril orientation. (B2-H2) Images were post-processed with orientation-J plugin in ImageJ to build a colour map of fibril orientation.

## 3. DISCUSSION

Hybrid biofabrication processes can leverage the specific advantages of different additive manufacturing (AM) technologies [30], creating new and otherwise inaccessible opportunities to expand fundamental concepts of tissue engineering. Here, we hypothesised that the process of cartilage self-assembly could benefit from the association of MEW and inkjet bioprinting. To this end, the objective was to jet cells into the individual chambers of MEW sheets with a view to driving cellular condensation and cartilage-specific tissue organization. We demonstrated that the combination of MEW and inkjet printing supports the self-assembly of organized arrays of mesenchymal aggregates that fused to form a highly connected tissue with sGAG content approaching that of native cartilage. The polymeric chambers were able to drive an articular cartilage-like histotypical organization within the hybrid tissue; specifically, the cell and collagen fibrils organization were found to match that of skeletally immature joints. It was also possible to integrate inkjet printing and MEW to engineer clinically sized cartilage grafts with biomechanical properties close to that of native cartilage. Taken together, these results demonstrate how the integration of different 3D-printing techniques can make it possible to produce functional stratified cartilage tissues with low-polymer content (< 2%) for joint resurfacing applications.

The combination of MEW hydrophobic PCL fibers and non-adhesive coating provided an environment that supported cellular condensation within each microchamber, interactions between cell-aggregates, and their functional development. In a previous study, we used FDM to print a polymeric framework on a non-porous PCL base where spheroids self-assembled and produced robust stratified cartilage [20]. However, the piling of thick PCL fibers prevents cell-cell communication, whereas the stacking of small MEW fibers with random-opened frames in the fiber wall (due to more fibers at the interconnections than on the wall itself) offers multiple opportunities for aggregates to physically connect, even at an early stage, and generate highly dense tissues. In addition, the use of non-adhesive coating as a temporary base offers the possibility to easily remove the engineered tissue from the culture insert and use it as a patch for joint resurfacing, which would not be possible with a solid base fused to the microchamber system. The highly compliant nature of the MEW sheets also allows them to be fitted over complex surfaces post-printing (**Supplementary Figure 1.F**), which would not be possible using more rigid polymeric structures generated using additive manufacturing techniques such as FDM.

It is clear from SEM and macroscopic pictures that cells were initially localised in the interior of the microchambers, and the resulting aggregates grew over time in culture and fused with adjacent spheroids through and over the surface of the microchambers. The engineered tissues had an sGAG content approaching that observed in native cartilage after only 21 days in culture [18]. Hybrid constructs were also hyaline-like in composition, staining strongly for type II collagen and weakly for type I collagen. While total collagen content at this timepoint was less than that of native tissue, the organization of the collagen network mimicked that of articular cartilage taken from 12 weeks old porcine synovial joints, with a three-layer organization typical of juvenile cartilage [55,56]. It should be noted that the colour and intensity of collagen birefringence are influenced by the alignment of the collagen fibrils, the packing density [57], the diameter of the fibrils [58], and the presence of proteoglycans in the tissue section [59]. Hence, densely packed and highly aligned collagen fibrils of the tangential and isotropic zones display a yellow birefringence, whereas the loosely woven and developing radial zone is seen with a green birefringence, which will turn yellow as fibril diameter and alignment will increase with aging [18,55]. This is well supported by HIM observations revealing a stratified collagen network composed of thin collagen fibrils and becoming denser toward the surface of the tissue. The fact that cells can reorganize extracellular matrix (ECM) components during collective migration is well known [30], and it has been shown that PCL microfibers can guide the growth of collagenous tissues *in vivo* [60]. Furthermore, it is also known that achieving some level of structural organization in the collagen network is possible by spatial-confining self-assembling tissues [15]. Therefore, this structured organization observed in hybrid tissues is likely due to the orientated growth enforced by the boundaries of the MEW fibers. This anisotropic organisation will help the implant withstand immediate load bearing and is anticipated to mature into the classical Bennighoff architecture observed in mature cartilage following *in vivo* implantation. While the overall collagen content was found below that of native tissue, the MEW fibers will also function to provide tensile strength and stiffness to the engineered tissue while the secreted collagen network continues to mature following implantation.

The utility of the newly developed biofabrication process was further demonstrated by generating self-assembled spheroids in a scale-up cartilage implant composed of ≈ 3,000 microchambers. Such control would not have been possible without using biofabrication technology. The deposition of articular cartilage ECM overtime led to a dramatic increase in compressive properties, approaching that of skeletally immature articular cartilage [18]. The dynamic modulus, which is correlated with the integrity of the tissue’s collagen network and secondarily to the swelling pressure induced by proteoglycans [50], was in the MPa range. MEW scaffolds have also previously been used to mechanically reinforce soft hydrogels [31,61]. These studies showed that the reinforcement effects during compressive loading were associated with the resistance of the fibre cross-section interconnections [61], and fibres being put under tension by lateral hydrogel expansion [31]. The reinforcement to compressive loading observed here is likely due to a synergistic interaction between the MEW fibers and the deposited ECM, producing a composite with time dependant mechanical properties similar to native tissue [62], together with the developing Benninghoff architecture that also improved the mechanical properties. It is anticipated that this mechanical reinforcement would help the implant withstand immediate load-bearing in a joint environment.

While the compressive mechanical properties of the hybrid tissue were dramatically higher than that of empty control, the MEW network still dominated the tensile properties of the construct after 8 weeks of *in vitro* maturation. Previous studies have shown that the collagen matrix buckles under compressive strains in proteoglycan-depleted cartilage [23], pointing to its primarily role in providing tensile strength and stiffness in the tissue [19]. Indeed the compressive modulus of such proteoglycan-depleted cartilage can be as low as 3 kPa, which is comparable to our empty MEW scaffolds. To the best of our knowledge, no tissue engineering strategy has to date been able to generate constructs with native-tissue like levels of collagen content and organization. We believe that the relatively low collagen content observed in our engineered tissues, coupled with a collagen network organization more mimetic of juvenile articular cartilage than fully mature cartilage, could explain the finding that the matrix deposited within the MEW network did not significantly increase the overall tensile modulus of the graft. However, *in vitro* maturation did increase the tensile yield strength of the construct, and the tensile ramp modulus of hybrid tissues was still in the MPa range. It is anticipated that the maturation and reinforcement of the collagen network after implantation will improve the tensile properties, like in postnatal development [63]. Lastly, the tensile modulus of hybrid tissues was higher than the compressive modulus (tension-compression non-linearity), which is typical of articular cartilage mechanical properties and plays a fundamental role in its ability to support physiological levels of stress [48,64]. Our results suggest that the small PCL fibers printed by MEW played a key role in achieving non-linear tension-compression behaviour by directing the growth and maturation of self-assembled MSCs aggregates into cartilage tissue and secondly by providing tensile reinforcement.

Tissue engineering scalable cartilage grafts requires satisfying the physio-chemical demands of large volumes of cells, which can lead to inhomogeneous deposition of cartilage matrix within engineered tissues if nutrient and oxygen are not sufficiently provided during *in vitro* maturation [65–67]. These considerations become exacerbated with scaffold-free approaches which require high cell-seeding density to engineer even small tissues. Here, the culture of hybrid tissues was performed in static conditions, which may have led to areas with insufficient nutrients for spheroid maturation and fusion, and hence tissues with non-anatomically relevant thickness. Introducing dynamic bioreactor culture should address these points and increase the overall mechanical properties and thickness of the engineered grafts [20]. Future work is also required to evaluate if the hybrid tissues can integrate, sustain relevant mechanical loading, and further mature *in vivo*. Although we hypothesized that this new biofabrication strategy could be used to resurface complex joints, we did not address this challenge. Future work will look into the possibility to combine inkjet bioprinting with anatomically relevant scaffolds printed by MEW [29] for complex joint resurfacing. For example, microchambers could be printed directly onto a curved surface [68], and stratified cartilage engineered by self-assembling onto complex orthopedic implants such as a hip implant.

## 4. CONCLUSION

In this experimental work, we combined different AM technologies (MEW, inkjet) to engineer stratified cartilage tissues. The majority of bioprinting approaches developed to engineer cartilage have used overly-stiff and non-compliant structures to reinforce hybrid tissues, which does not mimic the role of the collagen network in articular cartilage. In addition, recapitulating the stratified zonal architecture of this tissue is a major challenge in the field. We have addressed both these challenges by generating arrays of spheroids within a MEW polymeric framework that orientated the growth and organization of the ECM secreted by the cells. This resulted in scalable tissues with a spatial collagen fibre organization mimicking that of skeletally immature joints and exhibiting tension-compression non-linear behaviour. Overall, this new hybrid biofabrication approach can create more biomimetic hybrid structures over existing methods, and could represent an alternative option for the treatment of cartilage defects.

## 5. MATERIALS AND METHODS

### Cell isolation and expansion

Bone marrow-derived mesenchymal stem cells were isolated from the femoral shaft of a porcine donor (Danish Duroc, male, 4 months old). The extracted marrow was washed in expansion medium containing high-glucose Dulbecco’s Modified Eagle Medium (hgDMEM), 10% foetal bovine serum (FBS) and penicillin (100 U/mL) – streptomycin (100 μg/mL) (all from Bioscience) and triturated with a 16G needle until a homogenous mixture was obtained. The suspension was then centrifuged at 650g for 5 min and the resultant cell pellet resuspended in fresh expansion medium twice before it was filtered through a 40 μm cell sieve (Sarstedt). Cell counting was performed with trypan blue in the presence of acetic acid (6% final) before plating at a density of 1.3 × 10^5^ cells/cm^2^. Following colony formation, cells were trypsinized, counted, and re-plated for 2 additional passages at a density of 5 × 10^3^ cells/cm^2^ at 5% pO2 in expansion medium supplemented with 5 ng/ml of fibroblast growth factor (FGF)-2 (PeproTech Ltd). Medium change was performed three times per week.

### Biofabrication process (melt-electrowriting, poly-HEMA coating and inkjet-bioprinting)

All constructs were printed with the 3D Discovery multi-head printing system (RegenHu, Switzerland). Melt-electrowriting (MEW) was performed with polycaprolactone (PCL, Capa® 6500D, Perstorp UK Ltd) molten in a metallic cartridge at 100 °C. PCL was extruded through a 24G nozzle with an air pressure of 0.06-0.08 MPa and voltage of 10 kV. The printhead was kept at a constant Z-coordinate of 3 mm and translated at a speed of 40 mm/sec in × and Y directions over a fixed collector plate. The MEW jet was stabilized before printing by printing 8 lines which were analysed for deviations in fibre diameter and/or pulsing. 60 × 60 mm box-like structures composed of 200 layers were printed. Each fibrous layer was orientated at 90° to the previous layer with 0.8 mm spacing between fibres. Accordingly, the walls of the pores consisted of 100 stacked fibers. The scaffolds were subsequently cut into 8 × 8 mm squares with a scalpel or directly sterilized with ethylene oxide.

To prevent cell adhesion, 12 well plates or 60 mm Petri dishes (Corning) were coated with 1,2% (w/v) Poly(2-hydroxyethyl methacrylate) (poly-HEMA, Sigma-Aldrich, ref. P3932) at a density of 70 μl/cm^2^ as previously described [69]. Sterile MEW scaffolds were placed onto the poly-HEMA coating and kept in place with a custom made metal or PCL ring so that the inkjet area was a 6 × 6 mm or a 50 × 50 mm square. Scaffolds were carefully washed with 1x Phosphate Buffered Saline (PBS) solution 3 × 5 min to set connections between the two hydrophobic materials (poly-HEMA coating and PCL) and to prevent the scaffolds from being resuspended when culture medium was added after cells were inkjet.

For inkjet bioprinting, a piezoelectric valve with an inner diameter of 0.3 mm and a stroke of 0.1 mm (Fritz Gyger AG, Switzerland, ref. 00015815) was attached to a contactless dispensing printhead. The printhead was aligned with the centre of a single microchamber and was kept at a constant Z-coordinate of 40 mm. Next, the printhead was translated according to an alternating horizontal path at a speed of 4 mm/sec. The piezoelectric valve opened for 1300 μsec every 0.8 mm to inkjet cells resuspended in expansion medium at a density of 30 × 10^6^ cells/ml with an air pressure of 0.1 MPa. Post-printing, constructs were placed for 10 min in the incubator to initiate cell aggregation before adding excess expansion medium to the construct.

### Chondrogenic conversion

Chondrogenic medium was added to the constructs 48 hours after inkjet bioprinting and consisted of hgDMEM supplemented with penicillin (100 U/mL) – streptomycin (100 μg/mL), 100 μg/ml sodium pyruvate, 40 μg/ml L-proline, 50 μg/ml L-ascorbic acid-2-phosphate, 4.7 μg/ml linoleic acid, 1.5 mg/ml bovine serum albumin (BSA), 1 × insulin–transferrin–selenium, 100 nM dexamethasone (all from Sigma-Aldrich), and 10 ng/ml human transforming growth factor-beta (TGF-b) 3 (PeproTech Ltd). Cells were cultured at 5% pO2 for at least 21 days and up to 8 weeks with medium change performed every two days

### Scaffold imaging and spheroid measurement

Cell-seeded scaffolds were imaged with an Olympus SZXY stereomicroscope and an Olympus IX71 optical microscope. To measure the size of spheroids, the freehand selection tool in ImageJ software (National Institutes of Health, USA) was used to measure the area of the cell aggregate. The diameter of a circle of equal projection area (Da) was then calculated using the equation Da = 2√(A/π) where A is the area measured [70].

### Scanning electron microscopy

Samples were fixed in 3% glutaraldehyde in 0.1 M cacodylate buffer (all from Sigma-Aldrich) at 4°C for a minimum of 12h. They were then rinsed twice in 0.1 M cacodylate buffer for 10 min, dehydrated in graded ethanol baths series, immersed twice in hexamethyldisilazane (Sigma-Aldrich) for 30 min and allowed to completely dry overnight before imaging. Samples were imaged with a Zeiss ULTRA plus scanning electron microscope and imaged coloured with GIMP software (version 2.10.22).

### Live/Dead confocal microscopy

Cell viability was assessed using a Live/Dead assay kit (Bioscience). Tissue constructs were rinsed with PBS and incubated in PBS containing 4 μM ethidium homodimer-1 and 2 μM calcein for 30 min. Samples were rinsed again in PBS and imaged with a Leica SP8 scanning confocal microscope at 515 and 615 nm channels. Images were analysed using Leica Application Suite × (LAS X). All images presented are 3D Z-stack reconstructions of the tissue. The depth reconstruction images were produced with Imaris software (BITPLANE, Oxford Instruments).

### Time-lapse cell imaging

The self-assembling of spheroids was imaged with a CytoSMART™ Lux2 system (CytoSMART technologies, Netherlands).

### Biochemical analyses

After 21 days of in *vitro culture* constructs were washed in PBS, weighed, and frozen for subsequent analyses. Each construct was digested with papain (3.88 units/ml) in 100 mM sodium phosphate - 5 mM ethylenediaminetetraacetic acid (EDTA) buffer (pH 6.5) with 10 mM L-cysteine–hydrochloride (all from Sigma-Aldrich) at 60 °C and 10 rpm for 18 hours. DNA content was quantified using the Hoechst Bisbenzimide 33258 dye assay, with a calf thymus DNA standard. The amount of sulphated glycosaminoglycan (sGAG) was quantified using the dimethylmethylene blue dye-binding assay (DMMB) (Blyscan, Biocolor Ltd.), with a chondroitin sulphate standard. Total collagen content was determined by measuring the hydroxyproline content using the dimethylaminobenzaldehyde and chloramine T assay, and a hydroxyproline to collagen ratio of 1:7.69 [71]. The weight of PCL was excluded from all tissue wet weight (ww%) normalisations.

### Histological and immunohistochemical analyses

Engineered tissue constructs were fixed in 4% paraformaldehyde, dehydrated in a graded series of ethanol’s, embedded in paraffin wax and sectioned at 5 μm. The sections were stained with hematoxylin and eosin to study cell morphology, alcian blue to reveal the presence of sGAG and picrosirius red to visualize the collagen content.

Collagen types I and II were also evaluated using a standard immunohistochemical technique. Rehydrated sections were treated with pronase (32 PUK/ml, Sigma-Aldrich) at 37°C for 5 min, then incubated in blocking buffer containing 1% (w/v) BSA and 10% (v/v) goat serum (all from Sigma-Aldrich) in 1X PBS for 1 hour at room temperature (RT) to block non-specific sites. Tissue sections were then incubated with type I collagen (Abcam, ref. 90395, mouse monoclonal IgG, 1:400) or type II collagen (Santa-Cruz, ref. sc-52658, mouse monoclonal IgG, 1:200) primary antibody diluted in blocking buffer overnight at 4°C in a humidified chamber. Samples were then incubated with 3% (v/v) hydrogen peroxide solution (Sigma-Aldrich) for 20 min to block endo-peroxydase activity, then with secondary antibody (Sigma-Aldrich, ref. B7151, anti-Mouse IgG) diluted in blocking solution (1.5:200 for detection of type I collagen and 1:300 for detection of type II collagen) for 1 hour at RT. Following a 45 min incubation period with ABC reagent (ABC Elite kit Vectastain PK-400, Vector Labs), the DAB substrate (SK-4100, Vector Labs) was added to the tissue section and the presence of the protein of interest was revealed by the apparition of brown staining in the positive control. Histological and immunohistochemical samples were imaged with a slide scanner (Scanscope, Leica biosystems) and analysed with the Aperio software (Leica biosystems).

### Polarized light microscopy

Rehydrated tissue sections were incubated at 37°C for 18 hours with 1000 U/ml bovine testicular hyaluronidase (Sigma-Aldrich, ref. H3506) prepared in 0.1M phosphate buffer pH 6.9 to remove proteoglycans so birefringence was only caused by collagen fibrils [72,73]. Sections were then stained with 0.1% (w/v) picrosirius red, mounted with DPX (all from Sigma-Aldrich) and imaged with an Olympus BX41 polarizing light microscope equipped with a MicroPublisher 6™ CCD camera and an Olympus U-CMAD3 adaptor. Average orientation, dispersion and coherency of collagen fibrils in the engineered tissue and articular cartilage (12 weeks old pig, Danish Duroc, control sample) were assessed using orientation-J and directionality plugins in ImageJ [74].

### Helium ion microscopy

Engineered tissues were imaged with a helium ion microscope (Zeiss ORION Nanofab) for high-resolution visualization of the collagen network. Before imaging, serial enzymatic digestion was used to remove glycosaminoglycan to provide an unobstructed view of the collagen fibrils; this protocol is based on the method described by Vanden Berg-Foels *et al*. [75]. Images were acquired in secondary electron mode with an acceleration voltage of 29.9 kV, a beam current of 1.54-1.72 pA, and a dwell time of 2-5 μs. Images were acquired using a pixel resolution of 1024 × 1024 or 2048 × 2048. The brightness and contrast were optimised and images were analysed with orientation-J plugin in ImageJ; no other post-processing procedures were performed.

### Mechanical testing

Unconfined compression tests were carried out on samples produced in the shape of a cylinder using a 5 mm diameter biopsy punch. Samples were placed in a PBS bath at room temperature (∼ 25 °C) and compressed using a twin column Zwick universal testing machine (Zwick, Roell) equipped with a 100 N load cell. A preload of 0.02 N was used for empty scaffolds, whereas 0.5 N preload was applied to the tissue-engineered cartilage. A combined stress-relaxation and dynamic compression protocol was applied in increasing steps of 10% to a maximum of 30% [76]. Peak strain was reached within 500 seconds followed by 45 min relaxation. Five compressive cycles at 1% strain and 1Hz frequency were then superimposed. The compressive (or ramp) modulus was taken as the slope of the stress-strain curve between 10%-20% and 20%-30% strain. The equilibrium modulus was determined for the last 10 seconds of the equilibrium phase following unconfined compression testing to 20% and 30% strain. The dynamic modulus was calculated from the average force amplitude over the five compression cycles following the relaxation test [42].

Stress-relaxation tensile tests for both the tissue-engineered constructs and PCL scaffolds were conducted on a TA Instruments TestBench with a 20 N load cell. The tensile samples were cut with a length to width ratio of 4:1 in the gauge section (20 mm × 5 mm) and an additional 5 mm for either grip section. The tests were characterized by an initial preload of 0.02 N and a sequential ramp through three relaxation points at 3, 6, and 9% strain. These points were shown to be below the yield point *via* uniaxial tensile testing. For each phase, a constant ramp rate of 0.3 mm/min and an equilibrium time of 30 min were used. After the final relaxation point, the test was ramped beyond the expected yield point to 40% strain. Throughout the test, samples were kept hydrated via PBS drips. Both the ramp and equilibrium tensile modulus were calculated for each relaxation phase. The ramp modulus was calculated as the slope of the stress-strain curve for linear regions approaching each equilibrium point. The equilibrium modulus was assessed by averaging the last 10 force readings from the load cell of each equilibrium phase. Finally, the yield point was calculated *via* an offset from the initial ramp modulus at 0.2% strain.

### Statistical analyses

Statistical analyses were performed using the software package GraphPad Prism (Version 7.00). Statistical tests used to assess the normal distribution of data or to compare groups are indicated in figure legends. When groups were compared, significance was accepted at a level of p < 0.05. Results are expressed as mean ± standard deviation. Graphical results were produced with GraphPad Prism.

## ACKNOWLEDGEMENTS

This publication was developed with the financial support of Science Foundation Ireland (SFI) under grant number 12/RC/2278 and 17/SP/4721. This research is co-funded by the European Regional Development Fund and SFI under Ireland’s European Structural and Investment Fund. This research has been co-funded by Johnson & Johnson 3D Printing Innovation & Customer Solutions, Johnson & Johnson Services Inc. SEM was carried out at the Advanced Microscopy Laboratory (AML), Trinity College Dublin, Ireland. The AML is an SFI supported imaging and analysis centre, part of the CRANN Institute and affiliated to the AMBER centre. The authors would like to thank Dr. Gavin McManus for his support with confocal imaging and Imaris software, as well as Dr. Mark Ahearne for kindly providing the Cytosmart live-cell imaging system.

## AUTHOR CONTRIBUTIONS

Conceptualization: A.D., D.K.; Formal analysis: A.D., D.K., O.G.; Funding acquisition: D.K., O.G.; Investigation: A.D., X.B.G., C.O, K.E., S.V.E.; Methodology: A.D., D.K.; Project administration: D.K., O.G.; Supervision: A.D., D.K., O.G.; Validation: A.D., D.K, O.G.; Visualization: A.D.; Writing - original draft: A.D., D.K.; Writing - review & editing: A.D., D.K, O.G.; All authors contributed to writing the manuscript. Competing interests: The authors declare that they have no competing interests. All authors have given approval to the final version of the manuscript.

## DATA AVAILABILITY STATEMENT

The data presented in this study are available on request from the corresponding author.

## Notes

### Competing Interest Statement

The authors have declared no competing interest.

