## Supplemental Material for "Integrating melt-electrowriting and inkjet bioprinting for engineering structurally organized articular cartilage"

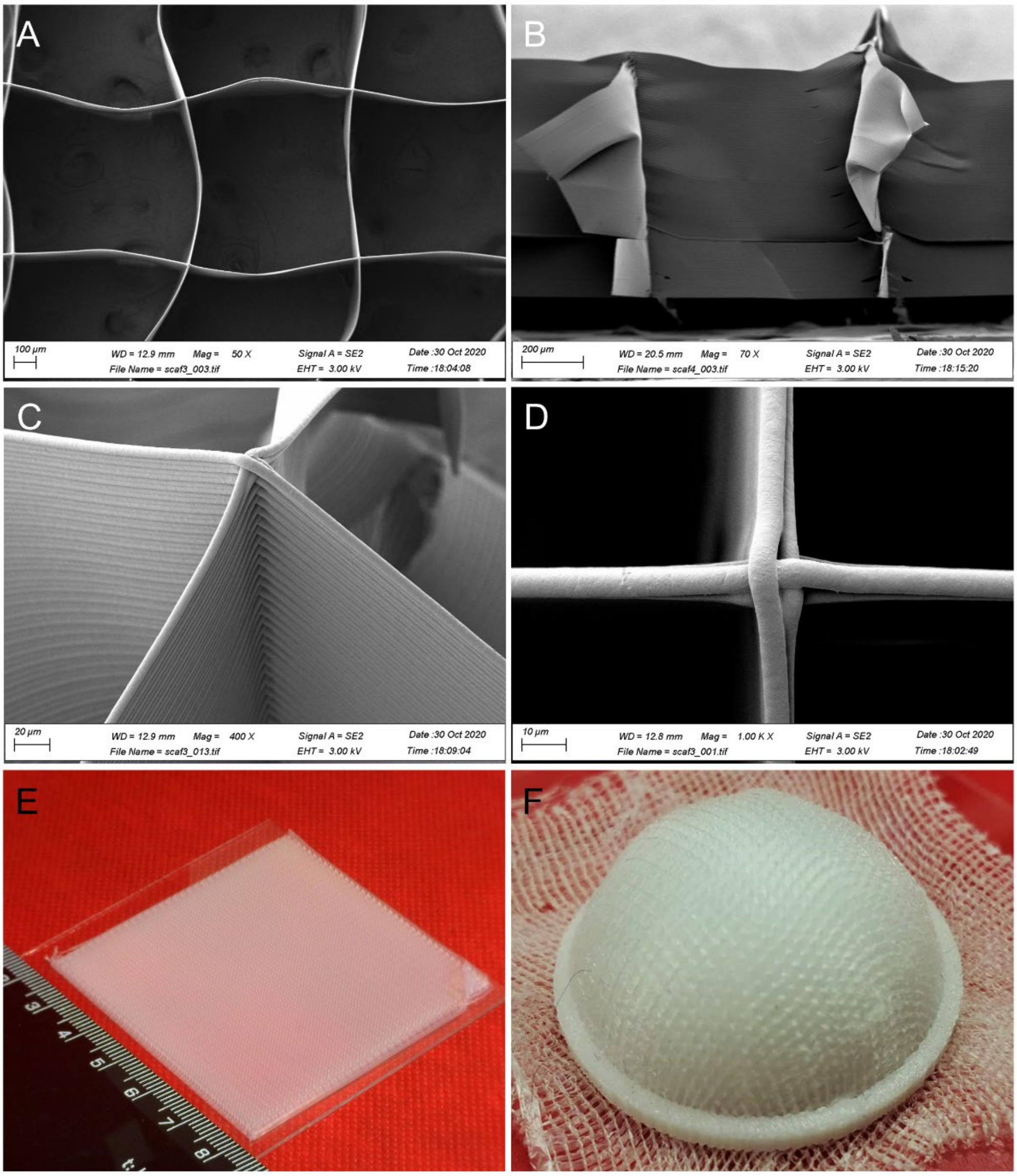


**SUPPLEMENTAL FIGURE 1: MEW scaffold.** (A-D) Scanning electron microscopy (SEM) and (E, F) macroscopic images of printed scaffolds. (E) Scale = cm. (F) The MEW scaffold was press-fitted onto a curved surface with a PCL ring.

| Collector speed [mm sec−1] | Pressure [MPa] | Box width [µm] | Wall fiber diameter [µm] | Wall fiber thickness [µm] | Voltage (KV) |
|---|---|---|---|---|---|
| 40 | 0.07 | 809 ± 46 | 7 ± 0.5 | 751 ± 42 | 10 |

**SUPPLEMENTAL TABLE 1**: **MEW parameters and corresponding scaffold dimensions determined from SEM images.**
